# Autophagy protects against high-dose *Mycobacterium tuberculosis* infection

**DOI:** 10.1101/2022.11.04.515158

**Authors:** Siwei Feng, E. Michael Nehls, Rachel L. Kinsella, Sthefany M. Chavez, Sumanta K. Naik, Samuel R. McKee, Neha Dubey, Amanda Samuels, Amanda Swain, Xiaoyan Cui, Skyler V. Hendrix, Reilly Woodson, Darren Kreamalmeyer, Asya Smirnov, Maxim N. Artyomov, Herbert W. Virgin, Ya-Ting Wang, Christina L. Stallings

## Abstract

Host autophagy had been associated with the control of *Mycobacterium tuberculosis* (Mtb) infection due to its ability to sequesters microorganisms through a process termed “xenophagy”^1–4^. Xenophagy purportedly limits Mtb replication within infected macrophages^1–4^. However, studies in mice using a standard low-dose infection model demonstrated that xenophagy in infected phagocytes is not required to control Mtb pathogenesis^5,6^. Instead, an autophagy-independent function of ATG5 in myeloid cells controls low-dose Mtb infection through limiting neutrophilic inflammation^5^. Hitherto, an *in vivo* role for autophagy during Mtb infection remained to be elucidated. We report herein that autophagy in myeloid cells mediates protection against high-dose Mtb infection, providing the first evidence for a role for autophagy in myeloid cells during Mtb infection *in vivo*. With the exception of ATG5, the autophagy proteins required to control high-dose Mtb infection are dispensable for host defense against a standard low-dose Mtb infection. Specifically, autophagy is required in CD11c^+^ cells, but is dispensable in neutrophils, to control a high-dose Mtb infection in the lung. The role for autophagy is not to directly degrade Mtb in macrophages through xenophagy, but mainly to limit myeloid-derived suppressor cell accumulation and to promote sustained protective T cell responses. Together, our data highlight a novel role for autophagy in controlling Mtb infection, distinct from that of Atg5 during low-dose Mtb infection, or any previously reported roles for autophagy. In addition, our finding that the result of a pathogen-plus-susceptibility gene interaction is dependent on pathogen burden has important implications on our understanding of how Mtb infection in humans can lead to a spectrum of outcomes, the variables that contribute to autophagy gene function during infection and inflammation, and the potential use of autophagy modulators in clinical medicine.

## Main Article

Pulmonary Mtb infection leads to either pathogen clearance, latent tuberculosis infection (LTBI), or active pulmonary tuberculosis (ATB) disease, the latter of which results in the clinical manifestations associated with TB. Although the infectious dose of Mtb in a given patient is unknown, work in animal models has demonstrated that higher infectious doses contribute to TB disease progression^7,8^ and different requirements for immune control^9–11^. Based on these studies, high-dose Mtb infection has been proposed to better recapitulate the infection outcomes and immunological responses observed in areas with high TB disease burden^12^. For wild-type C57Bl/6 mice, only a high dose (∼700-900 bacilli) of the Mtb HN878 strain, and not the standard low dose (∼100-450 bacilli) used in most studies, elicited gene expression profiles in the blood with similarities to the blood signatures observed in ATB patients^13^ These observations highlight that different immune mechanisms may be operative in the setting of different pathogen burden, however, our understanding of the immune requirements to control higher doses of Mtb infection remains incomplete^14^.

Aerosol delivery of high doses of Mtb (200-1000 colony forming units (CFU)) elicits progressive ATB in nonhuman primates (NHP), which is associated with a stronger macrophage interferon-γ (IFNγ) response compared to low-dose Mtb infection^7^. These elevated IFN responses resemble the elevated type I/II interferon signature in the blood of ATB patients^7,15,16^. BECLIN1 and FIP200, autophagy proteins required for initiation of autophagosome formation, mediate immune quiescence by dampening systemic IFNγ responses of tissue-resident macrophages^17^. As a result, mice lacking BECLIN1 in myeloid cells (*Becn1*^*f/f*^*-LysM-cre* mice) showed increased IFNγ responses and tissue resident macrophage activation across multiple organs, including in alveolar macrophages^17^. BECLIN1 and FIP200 do not share this role with other proteins that are essential for autophagy (i.e. ATG5, ATG16L1, and ATG7). Given the association between elevated macrophage IFNγ responses and TB disease progression during high-dose Mtb infection, we hypothesized that the role of BECLIN1 and FIP200 in dampening IFNγ responses of tissue-resident macrophages may impact Mtb pathogenesis in high burden scenarios, in a manner not observed in the low-dose Mtb infection model that is commonly used in mice. We infected *Becn1*^*f/f*^*-LysM-cre* mice with a high dose (∼1000 CFU) of Mtb strain Erdman and discovered that *Becn1*^*f/f*^*-LysM-cre* mice were extremely susceptible to high-dose Mtb infection, succumbing by 61 days post infection (dpi), while the majority of *Becn1*^*f/f*^ control mice survived passed 140 dpi (**Fig. 1a**). Deletion of other essential autophagy genes, *Atg14, Fip200, Atg7*, and *Atg16L1*, in LysM expressing cells also resulted in a similar susceptibility phenotype as observed in *Becn1*^*f/f*^*-LysM-cre* mice during high-dose Mtb infection (**Fig. 1b-1e**). Individual deletion of these essential autophagy genes, *Becn1, Atg14, Fip200, Atg7*, or *Atg16L1*, in mice has previously been shown to be sufficient to result in loss of autophagy in LysM^+^ cells^5,17,18^. In addition, since ATG7 and ATG16L1 function in later stages of autophagy than BECLIN1, these data indicate that the role of BECLIN1 in controlling high-dose Mtb infection is autophagy-dependent and distinct from its role in immune quiescence. These findings are also in contrast to data from low-dose (∼100 CFU) Mtb infection where *Atg5*^*f/f*^*-LysM-cre* mice exhibited increased susceptibility (**Fig. 1f**), a phenotype not shared by mice lacking other autophagy factors, including BECLIN1, ATG14, FIP200, ULK1/2, ATG4b, p62, ATG3, ATG12, ATG7 and ATG16L1^5,6^. We validated the previously published findings that *Atg14*^*f/f*^*-LysM-cre* are not more susceptible to low-dose Mtb infection, confirming that this essential role for ATG14 and the other autophagy proteins in controlling Mtb pathogenesis was unique to the higher dose infection (**Fig. 1g-h**).

**Figure 1.**
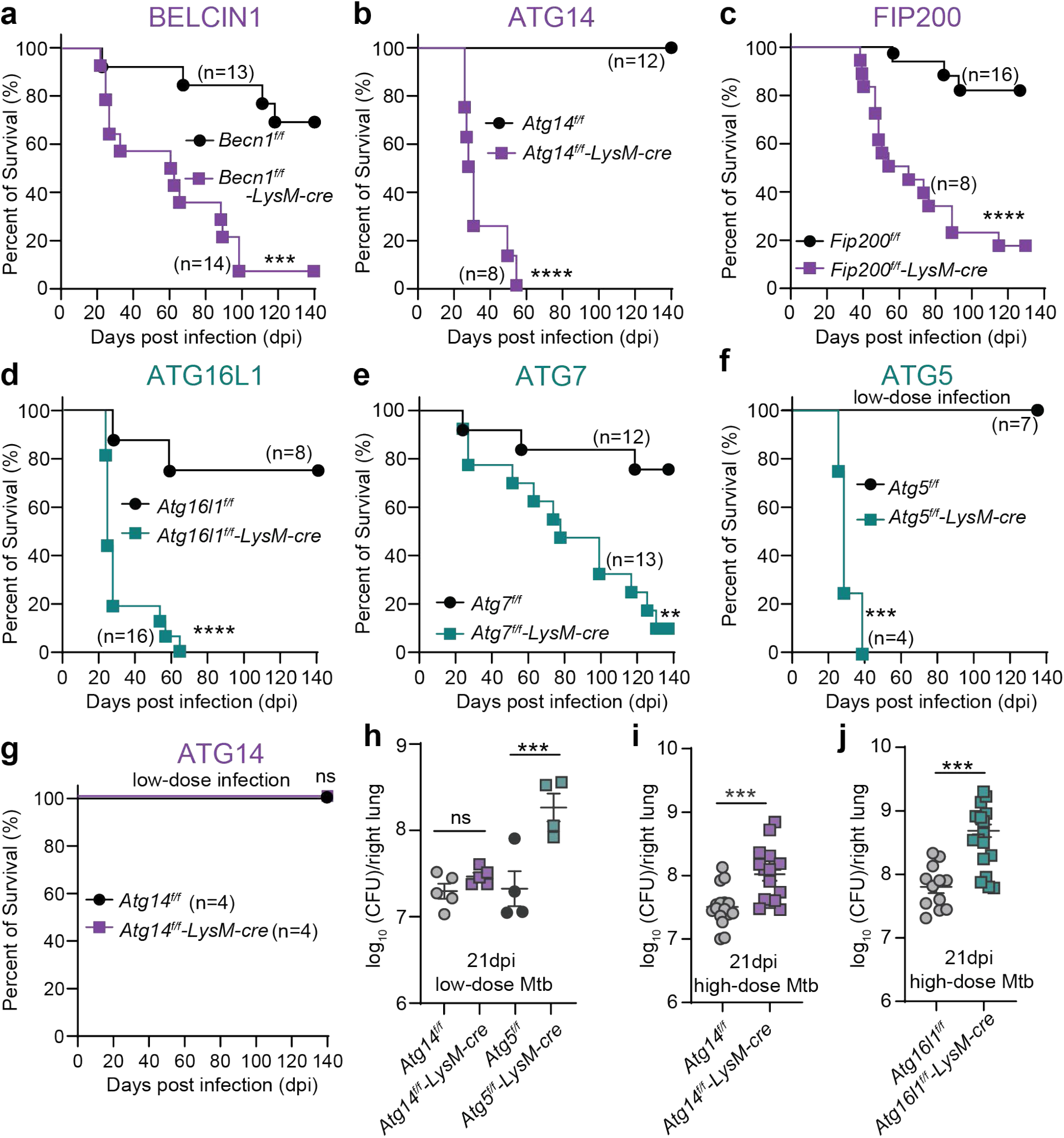
High-dose Mtb infection model uncovers protective role of autophagy in myeloid cells *in vivo*. **a-g**, Survival of mice that harbor myeloid deficiency (*LysM-cre*) in autophagy genes (*Becn1* (**a**), *Atg14* (**b, g**), *Fip200* (**c**), *Atg16l1* (**d**), *Atg7* (**e**), *Atg5* (**f**)) versus floxed littermate controls, after aerosol infection with high-dose of 1000 colony-forming units (CFU, **a**-**e**) or low-dose of 100 CFU (**f**-**g**) *M. tuberculosis* (Mtb) strain Erdman. *P* values were calculated using log-rank Mantel–Cox tests. **h-j**, Mtb CFU in lungs at 21 days post infection (dpi) of low-dose (**h**) and high-dose (**i-j**) infections. Means ± s.e.m. pooled from ⩾2 experiments are graphed. *P* values were calculated by two-tailed Mann-Whitney tests. ns= not significant. ** for *P* < 0.01, *** *P* < 0.001, and **** *P* < 0.0001.

To determine how autophagy genes in myeloid cells mediate protection during high-dose Mtb infection, we further characterized the most susceptible mouse lines: *Atg14*^*f/f*^*-LysM-cre* and *Atg16l1*^*f/f*^*-LysM-cre* mice (median survival time of 29.5 dpi and 24 dpi, respectively). The susceptibility of these mice to high-dose Mtb infection was associated with significantly higher lung bacterial burden at 21 dpi compared to control mice (**Fig. 1i-j**). These data support a protective role for multiple autophagy genes that is specifically revealed during high-dose Mtb infection.

We interrogated which myeloid cell type(s) require autophagy genes to control high-dose Mtb infection by comparing the susceptibility of mice that delete autophagy factors in LysM^+^ cells (neutrophils, macrophages, monocytes, and DCs), Mrp8^+^ cells (neutrophils), and CD11c^+^ cells (lung macrophages and DCs)^19,20^. Deleting *Becn1* or *Atg14* from neutrophils by Mrp8-cre^5^ resulted in wild-type level resistance (**Fig. 2a-b**), demonstrating that autophagy is not required in neutrophils during high-dose Mtb infection. Therefore, the role for autophagy during high-dose Mtb infection is genetically distinct from the autophagy-independent role for ATG5 in neutrophils during low-dose Mtb infection^4,5^. In contrast, loss of BECLIN1, ATG14, or ATG16L1 in CD11c^+^ cells resulted in increased susceptibility (**Fig. 2c-e**). Loss of ATG14 in CD11c^+^ cells also resulted in increased lung Mtb burden 21 dpi (**Fig. 2f**), similar to that observed in *Atg14*^*f/f*^*-LysM-cre* mice (**Fig. 1i**). These results indicate that multiple autophagy genes are individually required in CD11c^+^ macrophages and DCs to control high-dose Mtb infection, suggesting that autophagy as a cellular process, rather than the autophagy-independent role of ATG5 observed in the setting of low-dose challenge, is important in the high-dose infection model of TB.

**Figure 2.**
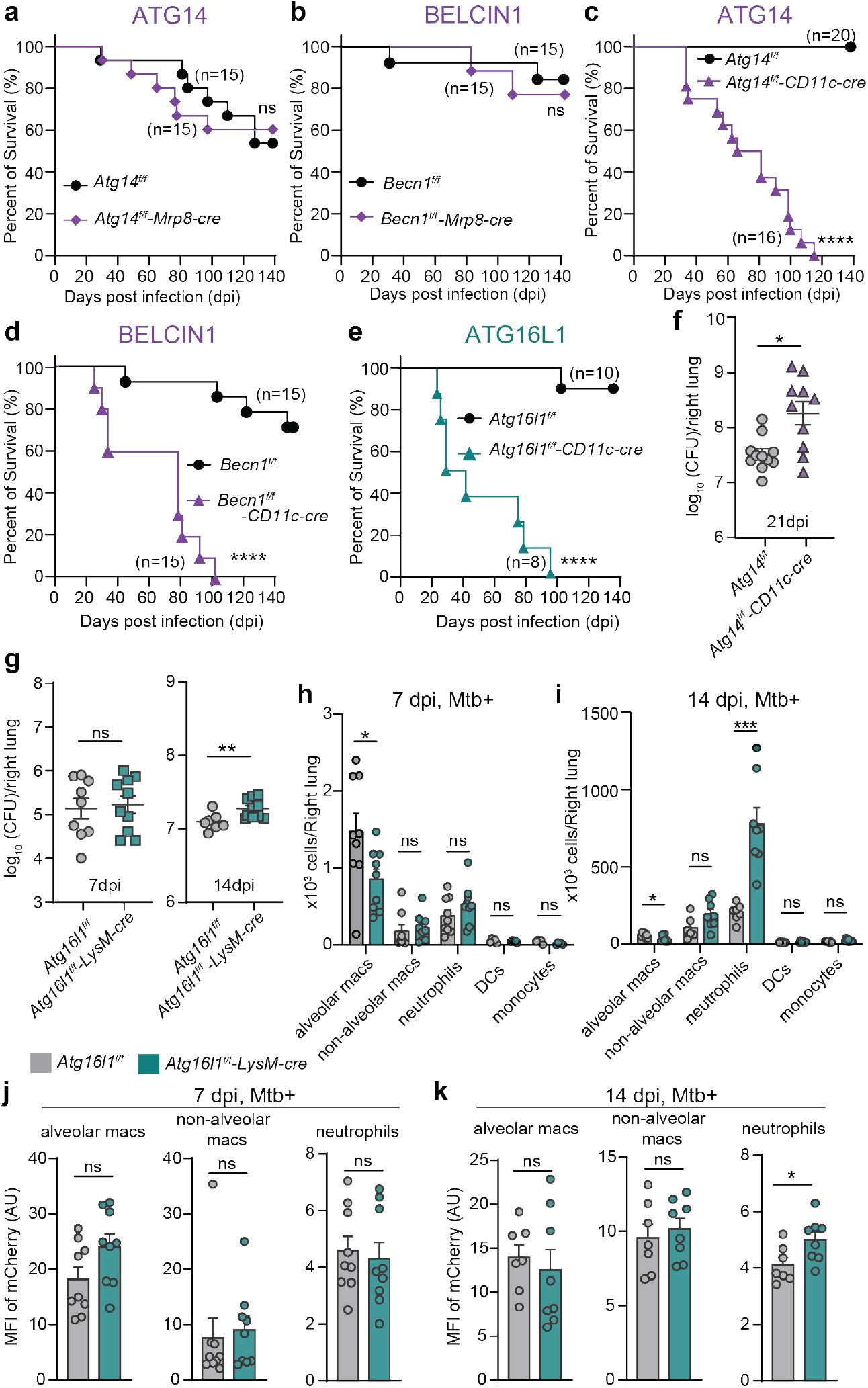
Autophagy genes in macrophages/DCs are required to control high-dose Mtb infection in mice. **a-e**, Survival of mice harboring *Becn1* (**a, c**), *Atg14* (**b, d**), *Atg16l1* (**e**) deletion in CD11c^+^ or Mrp8^+^ cells versus controls after aerosol infection with high-dose Mtb strain Erdman. *P* by log-rank Mantel–Cox tests. **f-g**, Mtb CFU in lungs at 21 dpi (**f**), 7 dpi or 14 dpi (**g**), with high-dose Mtb infection. Data are mean ± s.e.m., pooled from ≥2 experiments. *P* values were calculated using two-tailed Mann-Whitney tests. **h-k**, number of total mCherry+ cells in the left lungs at 7 dpi (**h**) and 14 dpi (**i**), and mCherry mean fluorescent intensity (MFI) in mCherry+ cells at 7 dpi (**j**) and 14 dpi (**k**), after high-dose infection with mCherry-Mtb Erdman. Data (Means ± s.e.m.) from 2 independent experiments are graphed. *P* values were calculated by two-tailed Mann-Whitney tests. *P* < 0.05 was denoted *, ** for *P* < 0.01, *** *P* < 0.001, and **** *P* < 0.0001.

In cell culture, loss of autophagy in macrophages results in increased Mtb replication by 1.5-3 fold^1,2,4,21,22^, which has been attributed to the inability of these macrophages to eliminate bacteria by xenophagy. These *in vitro* findings do not correlate to a role for xenophagy in lung macrophages *in vivo* during standard low-dose Mtb infection, where the increase in Mtb burden in *Atg5*^*f/f*^*-LysM-cre* mice results from increased neutrophil inflammation^5^. However, since Mtb burdens were higher in the lungs of mice deficient for autophagy in myeloid cells at 21 dpi (**Fig. 1i, 1j, 2f**), it is possible that xenophagy was required to control Mtb replication in macrophages during high-dose Mtb infection. Alveolar macrophages are the first cells infected upon inhalation of Mtb and are the initial cells wherein Mtb replicates^23,24^. The infected alveolar macrophages traffic to the interstitium, resulting in subsequent infection of interstitial macrophages and DCs as well as macrophages, monocytes, and granulocytes recruited from the blood^23^. Mtb CFUs in the lungs of *Atg16l1*^*f/f*^*-LysM-cre* mice at 7 dpi were similar to those in *Atg16l1*^*f/f*^ controls, suggesting that there is not an overall difference in bacterial replication early during infection (**Fig. 2g**). By 14 dpi, *Atg16l1*^*f/f*^*-LysM-cre* mice had higher Mtb lung burden (1.53 fold, *p*=0.0099) relative to control mice, indicating that ATG16L1 begins to control Mtb replication between 7 and 14 dpi (**Fig. 2g**). To determine what cell types harbored the replicating Mtb at 7 and 14 dpi, we infected mice with a high dose of an mCherry expressing Erdman strain of Mtb (**Fig. S1**). Similar to previous reports in wild-type mice^23,24^, the majority of cells infected at 7 dpi were alveolar macrophages (**Fig. 2h**) and the predominantly infected cell type at 14 dpi were neutrophils (**Fig. 2i**). The number of Mtb infected *Atg16l1*^*f/f*^*-LysM-cre* alveolar macrophages were lower than controls at both 7 dpi and 14 dpi (**Fig. 2h, 2i**). The levels of mCherry-Mtb per alveolar macrophage, measured by mean fluorescent intensity (MFI) of mCherry, at both 7 dpi and 14 dpi in *Atg16l1*^*f/f*^*-LysM-cre* mice were comparable to control mice (**Fig. 2j, 2k**). These data indicate that disabling autophagy genetically does not impact bactericidal activity of alveolar macrophages. Therefore, xenophagy is not required in alveolar macrophages to directly control bacterial replication early after infection. The other main cell types infected by high-dose Mtb were non-alveolar macrophages and neutrophils, which also harbored similar levels of Mtb in *Atg16l1*^*f/f*^*-LysM-cre* mice compared to *Atg16l1*^*f/f*^ controls at 7dpi (**Fig. 2h, 2j**). At 14 dpi, there were more Mtb infected neutrophils (**Fig. 2i**) and higher levels of Mtb per neutrophil in *Atg16l1*^*f/f*^*-LysM-cre* mice (**Fig. 2k**), which corresponds to when we detected higher CFUs in the lungs (**Fig. 2g**). These results indicate that the higher Mtb burdens in *Atg16l1*^*f/f*^*-LysM-cre* mice were due to higher levels of neutrophils being infected by 14 dpi, but not due to a defect in Mtb elimination in CD11c^+^ macrophages by xenophagy.

To determine how loss of autophagy in myeloid cells could impact Mtb infection and replication in neutrophils, we investigated neutrophil inflammation following high-dose Mtb infection. By 14 dpi, the frequency and number of neutrophils in *Atg16l1*^*f/f*^*-LysM-cre* lungs were greater than in the lungs of control mice (**Fig. 3a, S2**), in addition to the number of Mtb-infected neutrophils (**Fig. 2i, 3b**). These differences were more pronounced at 21 dpi (**Fig. 3a, 3b**). Consistent with the increased neutrophil inflammation, the lungs of *Atg16l1*^*f/f*^*-LysM-cre* mice at 14dpi and 21 dpi contained higher levels of proinflammatory cytokines, including G-CSF, M-CSF, IL-1α, IL-1β, IL-6, MIP-1b, MIP-2a, KC, and TNFα than control mice (**Fig. 3c**). GM-CSF, MCP-3, MCP-1, IL-18 and CXCL10 were also higher in *Atg16l1*^*f/f*^*-LysM-cre* lungs at 21 dpi, but not 14 dpi, when the level of neutrophil inflammation was highest in *Atg16l1*^*f/f*^*-LysM-cre* lungs (**Fig. 3a, 3c**). The higher numbers of neutrophils were associated with significant lower numbers of antigen presenting cells, alveolar macrophages and DCs, in *Atg16l1*^*f/f*^*-LysM-cre* mice at 21 dpi with high-dose Mtb, while numbers of non-alveolar macrophages and inflammatory monocytes were similar in the lungs of *Atg16l1*^*f/f*^*-LysM-cre* and *Atg16l1*^*f/f*^ mice (**Fig. 3d, S3**).

**Figure 3.**
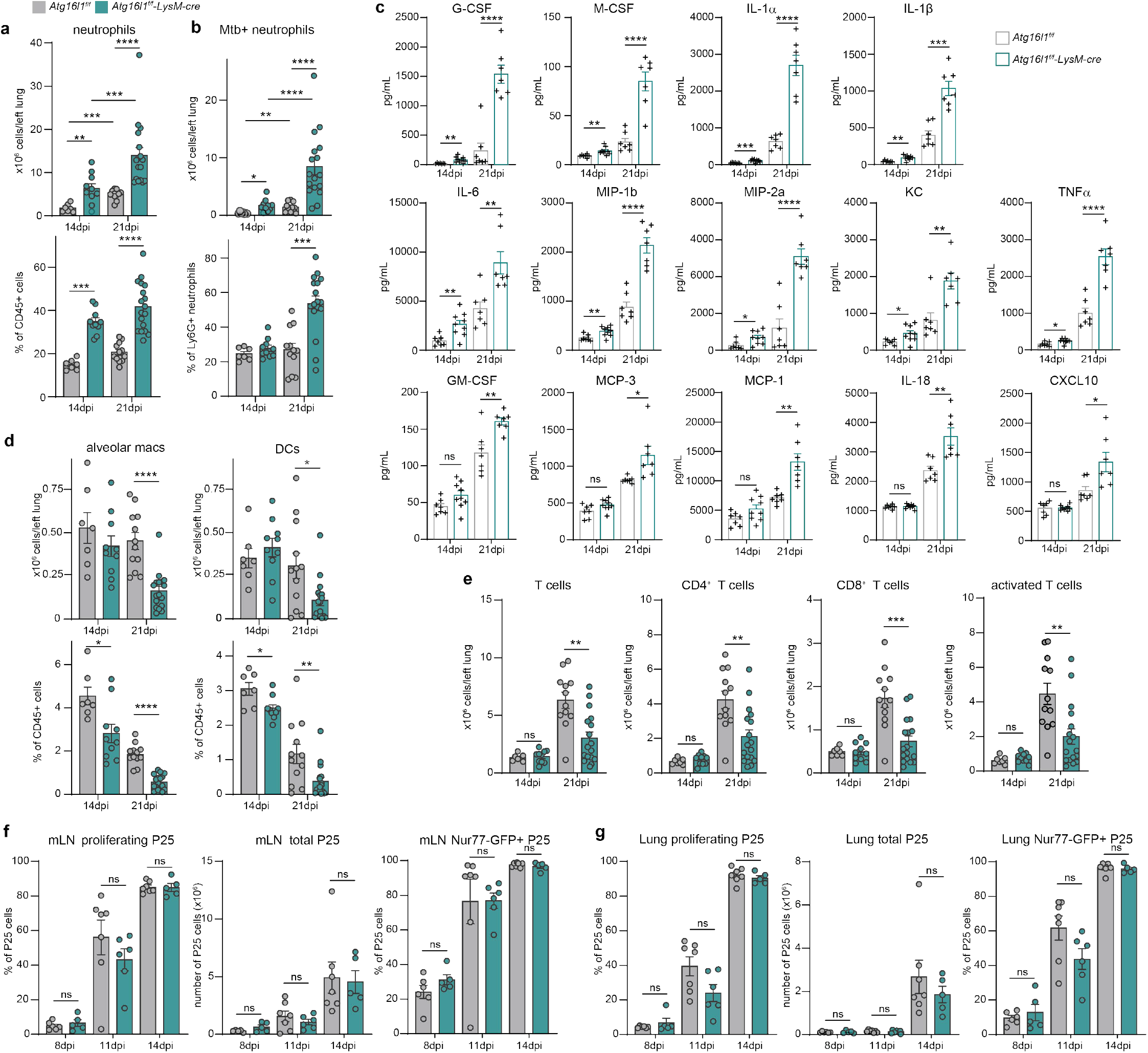
Myeloid autophagy suppresses neutrophil inflammation and promotes T cell response during high-dose Mtb infection. **a**,**b**,**d**,**e**, The number and percentage of immune cells in the lung of mice after high-dose Mtb infection. **c**, Concentration of cytokines in Mtb infected lungs as detected by multiplex cytokine panel, *P* values were calculated by two-tailed t-tests. **f**-**g**, The percentage of P25 CD4+ T cells that have undergone proliferation measured by CellTrace dilution, number of P25 CD4+ T cells, and the percentage of Nur77-GFP negative P25 CD4+ T cells in the mediastinal lymph nodes (mLN, **f**) and lungs (**g**) after high-dose Mtb infection. Means ± s.e.m. pooled from 2 independent experiments are graphed. *P* values were calculated by two-tailed Mann-Whitney tests. * for *P* < 0.05, ** for *P* < 0.01, *** *P* < 0.001, and **** *P* < 0.0001.

The lower numbers of alveolar macrophages and DCs prompted us to analyze the T cell responses in *Atg16l1*^*f/f*^*-LysM-cre* mice infected with a high dose of Mtb. We observed significantly fewer T cells and fewer activated T cells (CD44^hi^CD62L^lo^) in the lungs during high-dose Mtb infection in *Atg16l1*^*f/f*^*-LysM-cre* mice compared to controls at 21 dpi (**Fig. 3e, S4**), indicating a defect in T cell responses in these mice. After Mtb infection, antigen presenting cells migrate to the mediastinal lymph node (mLN) between 6-8 dpi to activate antigen specific T cells^25^. Mtb-specific T cells then proliferate and traffic to the lung to control Mtb replication^26^. To dissect whether the lower numbers of T cells at 21 dpi in the lungs of high-dose Mtb-infected *Atg16l1*^*f/f*^*-LysM-cre* mice was due to impaired T cell priming, we monitored the activation and proliferation of adoptively transferred Mtb-specific T cells (CellTrace labeled Ag85B specific CD4^+^ T cells^26^, referred to hereafter as P25 cells) in *Atg16*^*f/f*^*-LysM-cre* mice and control mice (**Fig. S5**). Proliferation of Mtb-specific P25 cells began between 8-11 dpi in the mLN after high dose Mtb infection of both *Atg16l1*^*f/f*^*-LysM-cre* mice and *Atg16l1*^*f/f*^ mice (**Fig. 3f**). The frequency and number of P25 cells proliferating in the mLN were similar in *Atg16l1*^*f/f*^*-LysM-cre* mice and *Atg16l1*^*f/f*^ mice at 11 dpi and 14 dpi (**Fig. 3f**). The number and frequency of proliferated P25 cells trafficked to the lung were similar in *Atg16l1*^*f/f*^*-LysM-cre* mice and *Atg16l1*^*f/f*^ mice at 11 dpi and 14 dpi (**Fig. 3g**). TCR-mediated cell activation, measured by Nur77-GFP positivity^27^, was also comparable in *Atg16l1*^*f/f*^*-LysM-cre* and *Atg16l1*^*f/f*^ mice between 8 and 14 dpi (**Fig. 3f-g**). Together, these analyses indicate that wild-type kinetics of T cell priming and arrival to the Mtb infected lung remain intact in the *Atg16l1*^*f/f*^*-LysM-cre* mice infected with a high dose of Mtb. Therefore, the lower T cell numbers in the lung at 21 dpi with high-dose Mtb in *Atg16l1*^*f/f*^*-LysM-cre* mice occurred after T cell priming and trafficking to the lung.

We noted that some of the neutrophils in the lungs of high-dose Mtb infected mice expressed an intermediate level of Ly6G and Gr-1 (Ly6G^int^ and Gr-1^int^) (**Fig. 4a, S6**), which resembles what has been described for myeloid derived suppressor cells (MDSC) that can potently suppress T cell function during cancer and chronic infections^28,29^. In addition, *Atg16l1*^*f/f*^*-LysM-cre* mice infected with high-dose Mtb accumulated a higher percentage of Ly6G^int^ cells at 14 dpi and 21 dpi compared to controls (**Fig. 4a**). Therefore, we hypothesized that the lower number of T cells in the *Atg16l1*^*f/f*^*-LysM-cre* mice during high-dose Mtb infection was due to increased neutrophil inflammation, including the accumulation of the Ly6G^int^Gr-1^int^ cells. To determine if depletion of neutrophils would improve T cell responses and subsequent control of Mtb, we depleted neutrophils by administering an anti-Ly6G (clone 1A8) antibody between 8-28 dpi using a dosing regimen previously shown to rescue *Atg5*^*f/f*^*-LysM-cre* mice as well as other susceptible mouse lines from low-dose Mtb infection^5,30^. Anti-Ly6G treatment in *Atg14*^*f/f*^*-LysM-cre, Atg16l1*^*f/f*^*-LysM-cre*, and control mice effectively eliminated Gr-1 high (Gr-1^hi^) neutrophils (**Fig. 4b**), but failed to provide any improvement in survival (**Fig. 4c, 4d**). Anti-Ly6G antibody treatment decreased Mtb burden in the lung of *Atg16l1*^*f/f*^*-LysM-cre* mice by 1.73 fold, and 1.43 fold in *Atg16l1*^*f/f*^ mice, compared to control-IgG treated mice (**Fig. 4e**), likely due to the decrease in Mtb infected Gr-1^hi^ neutrophils during 1A8 antibody treatment (**Fig. 4f**). However, anti-Ly6G antibody treatment did not affect the numbers of Gr-1^int^ neutrophils or the number of Mtb-infected Gr-1^int^ cells in the lungs of Mtb infected *Atg16l1*^*f/f*^*-LysM-cre* mice (**Fig. 4g, 4h**). Thus, the resistance of the Gr-1^int^ population to antibody-mediated depletion likely explains why the anti-Ly6G antibody treatment failed to completely revert the Mtb burden difference between *Atg16l1*^*f/f*^*-LysM-cre* and *Atg16l1*^*f/f*^ mice (**Fig. 4e**). Anti-Ly6G antibody treatment also failed to rescue the decrease in the numbers of alveolar macrophages, DCs, or T cells in *Atg16l1*^*f/f*^*-LysM-cre* lungs at 21 dpi (**Fig. 4i**), suggesting that accumulation of the Gr-1^int^ population could be contributing to the defects in antigen presenting cells and T cells.

**Figure 4.**
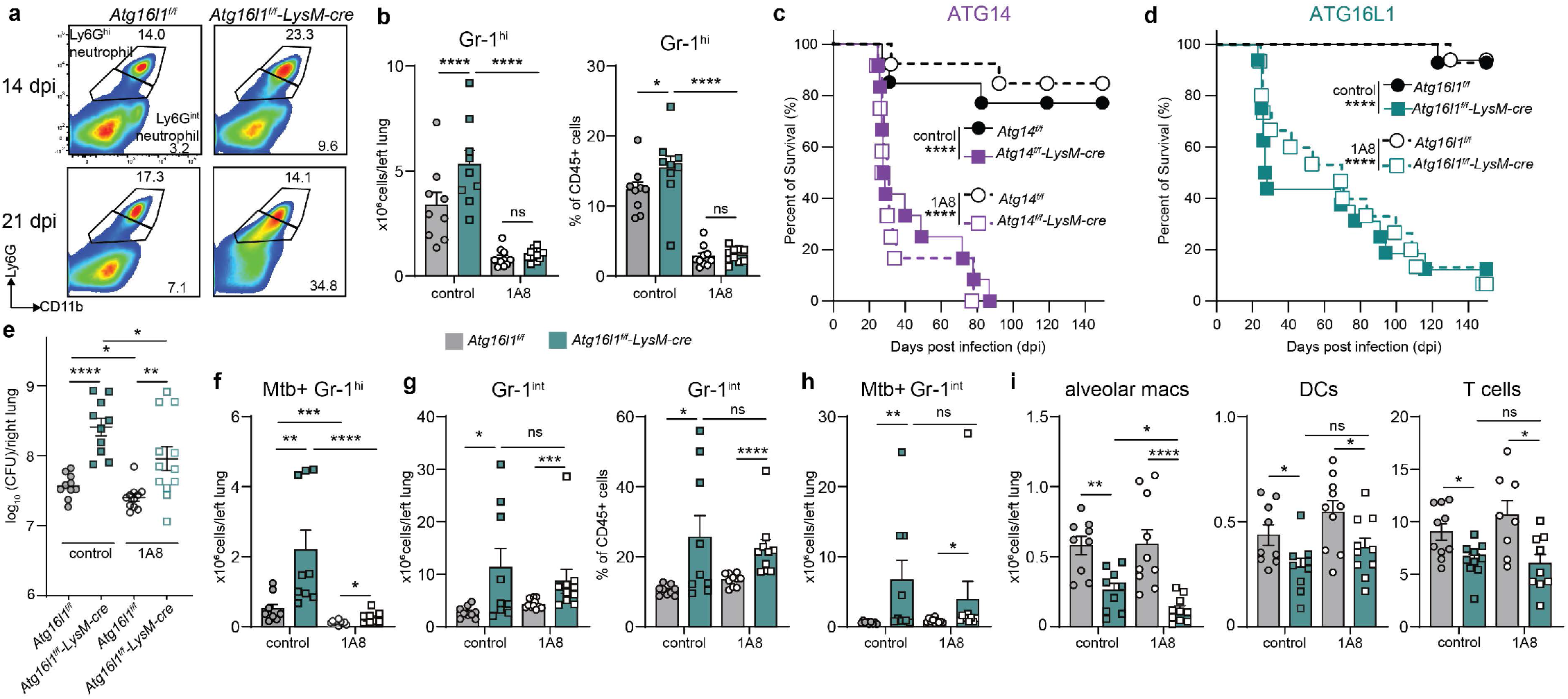
Accumulation of Gr-1^int^ neutrophils in autophagy deficient mice is associated with susceptibility and high burden. **a**. Representative flow cytometry plots of live CD45^+^MerTK^-^CD64^-^ cells from mice lungs. Data is representative of 3 mice from ≥3 experiments. **b**,**g**,**i**, The number and percentage of immune cells in high-dose Mtb infected mice and treated with neutrophil depletion antibody (1A8) or isotype control Ig every other day from 8-28 dpi. *P* values calculated by two-tailed Mann-Whitney tests. **c**-**d**, Survival of high-dose Mtb infected mice that were treated with 1A8 antibody or control immunoglobulin (control). *P* values calculated by log-rank Mantel–Cox tests. **e**, Mtb CFU in lungs 21 dpi. *P* values were calculated by two-tailed Mann-Whitney tests. **f**,**h**, Number of Mtb infected cells, measured as GFP+ cells in the lungs of mice 21 dpi of high-dose GFP-Mtb infection. *P* values were calculated by two-tailed Mann-Whitney tests. Data are mean ± s.e.m. pooled from 3 independent experiments. * for *P* < 0.05, ** for *P* < 0.01, *** *P* < 0.001, and **** *P* < 0.0001.

The Gr-1^int^ cells that accumulate in high-dose Mtb-infected *Atg16l1*^*f/f*^*-LysM-cre* mice are reminiscent of polymorphonuclear myeloid derived suppressor cells (PMN-MDSCs)^28,29,31^. To determine if the Gr-1^int^ neutrophils that accumulate during Mtb infection share characteristics with MDSCs, we performed single cell RNA-sequencing (scRNA-seq) on naïve and Mtb infected lungs from *Atg14*^*f/f*^, *Atg14*^*f/f*^*-LysM-cre* and *Atg14*^*f/f*^*-CD11c-cre* mice^5^. CD45^+^ clusters from lungs of naïve or 21 dpi of high-dose Mtb infected mice for each strain were projected into two dimensions with Uniform Manifold Approximation and Projection^32^ (UMAP) based on transcriptomic signature (**Fig. 5a**). Specific cell types were then stratified for unbiased sub-clustering. Within neutrophils, six *S100a8*^+^ sub-clusters (clusters PMN0-PMN5) were identified (**Fig. 5b**). Uninfected *Atg14*^*f/f*^*-LysM-cre* and *Atg14*^*f/f*^*-CD11c-cre* lungs showed altered neutrophil populations (**Fig. 5c**), which is consistent with the elevated basal lung inflammation previously reported for *Atg14*^*f/f*^*-LysM-cre* mice^33^. Mtb infection had a large effect on the neutrophil populations, where the naïve PMN0 and PMN5 populations were hardly detectable in the lungs at 21 days post high-dose infection (**Fig. 5c**). At 21 dpi, *Atg14* expression also affected the presence of specific neutrophil subsets, where PMN1 and PMN4 were the major clusters present in *Atg14*^*f/f*^*-LysM-cre* and *Atg14*^*f/f*^*-CD11c-cre* lungs, compared to PMN2 and PMN3 in *Atg14*^*f/f*^ (**Fig. 5c**). To identify signatures of pathways enriched in PMN1 and PMN4, we performed Gene Set Enrichment Analysis (GSEA) of a priori defined pathways and sets of PMN-MDSC signature genes identified in murine cancer models^34^ (**Fig. S7**). GSEA analysis demonstrated that compared to all PMN clusters, the most significantly enriched pathway in PMN1 was the activated-PMN-MDSC gene set from murine cancer (**Fig. 5d, S8a**). PMN4 was also distinctly enriched for the cancer PMN-MDSC gene set (**Fig. 5d, S8b**). Analysis of highly expressed genes in PMN1 and PMN4 revealed a substantial overlap in differentially overexpressed genes in PMN1 with activated-PMN-MDSC signature genes from murine cancer, and PMN4 with PMN-MDSC signature genes^15^ (**Fig. 5e**), indicating that PMN-MDSC-like cells elicited during Mtb infection in mice are highly similar to PMN-MDSCs in murine cancer models. Several genes upregulated in PMN1 and PMN4 were also enriched in human PMN-MDSCs from cancer patients (**Fig. 5e**). In addition, 11 highly enriched activated-PMN-MDSC and PMN-MDSC genes (*Pglyrp1, Plscr1, Gyg, Ifitm3, Ifitm1, Il1rn, Cd274, Gadd45b, Sod2, Npc2*, and *Hmox-1*) in PMN1 and PMN4 were among the signature transcripts within the blood of active TB patients^16,35^ (**Fig. 5e**), suggesting that the PMN-MDSC and activated-PMN-MDSC signatures are associated with worse outcomes in TB patients^8,36–40^.

**Figure 5.**
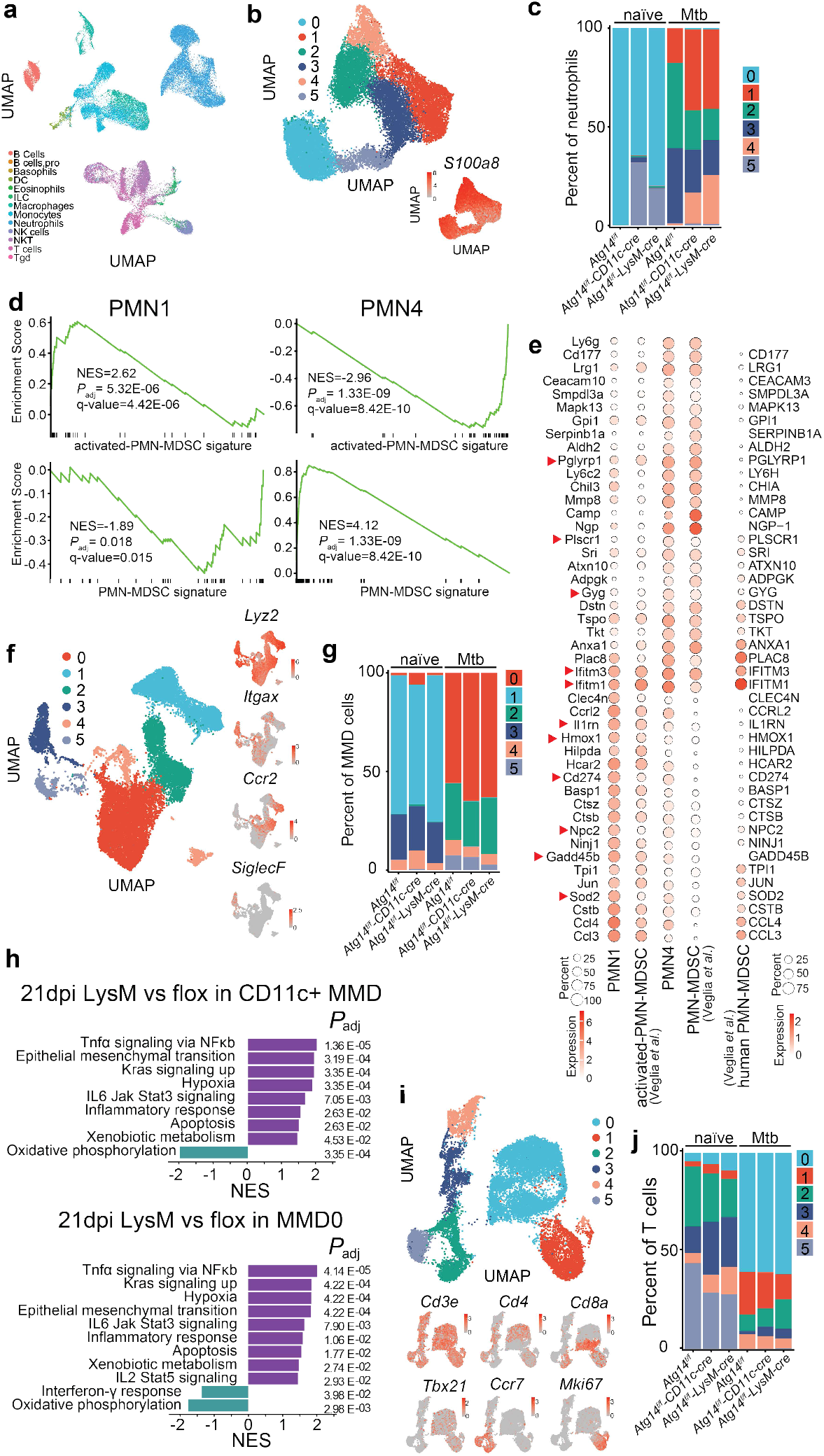
Autophagy deficiency in myeloid cells leads to accumulation of MDSC like cells during high-dose Mtb infection. **a**, UMAP plot of immune cells across all samples (lungs from naïve and 21 dpi of high-dose Mtb infected *Atg14*^*f/f*^, *Atg14*^*f/f*^*-CD11c-cre, Atg14*^*f/f*^*-LysM-cre* mice, n=4 samples per group, colored according to identified clusters. **b**,**f**,**i**, UMAP plots of neutrophils (**b**), macrophages/monocytes/DCs (MMD, **f**), and T cells (**i**) colored by clusters, and plots showing the expression level of markers genes. **c**,**g**,**j**, Distribution of neutrophil (**c**), MMD (**g**), and T cells (**j**) sub-clusters. **d**, Gene set enrichment analysis of activated-PMN-MDSC and PMN-MDSC signatures in neutrophil cluster 1 and 4 (PMN1, PMN4). The green curve represents the density of the genes identified in scRNA-seq analysis where normalized enrichment score (NES), *P*_adj_, and false discovery rate (q-value) are indicated. **e**, Dot plot showing highly expressed genes in PMN1 and PMN4, and expression of these genes in PMN-MDSCs of mouse cancer models, human cancer patients and active TB patients. Dot size represents prevalence of the transcript, and shade indicates expression level. The red arrows indicate genes identified among signature genes in active TB patients. **h**, significantly elevated and downregulated HALLMARK pathways found within all CD11c^+^ (*Itgax*) cells (top, see Fig. S9) and MMD0 (bottom) subcluster from infected *Atg14*^*f/f*^*-LysM-cre* lungs compared with infected controls as assessed by GSEA analysis.

The increased accumulation of MDSCs occurred in both *Atg14*^*f/f*^*-LysM-cre* and *Atg14*^*f/f*^*-CD11c-cre* mice during high-dose Mtb infection, supporting that autophagy genes function specifically in CD11c^+^ cells to suppress the accumulation of the PMN-MDSC populations (**Fig. 5c**). Therefore, we analyzed the gene expression profiles of the macrophages/monocytes/DCs (MMD) clusters to identify potential differences in these cell types that could be contributing to the MDSC accumulation during high-dose Mtb infection. The 6 MMD clusters identified were significantly shifted upon Mtb infection in all genotypes (**Fig. 5f-g**). Pathway analysis on the CD11c+ subclusters, which were defined by re-clustering MMD under higher resolution (**Fig. S9**), demonstrated that inflammatory pathways were upregulated in CD11c+ cells of *Atg14*^*f/f*^*-LysM-cre* and *Atg14*^*f/f*^*-CD11c-cre* mice at 21 dpi **(Fig. 5h, S10a)**, likely at least in part due to the higher bacterial burdens in these mice at that time point. In addition, apoptosis was among the significantly upregulated pathways in CD11c+ cells in *Atg14*^*f/f*^*-LysM-cre* mice after high-dose Mtb infection (**Fig. 5h**), consistent with the decreased macrophages and DCs observed in these mice at 21 dpi (**Fig. 3**). Specific analysis of the most prevalent CD11c+ cluster at 21 dpi, MMD0, revealed significant downregulation of the IFNγ signaling pathway in *Atg14*^*f/f*^*-LysM-cre* and *Atg14*^*f/f*^*-CD11c-cre* mice compared to infected *Atg14*^*f/f*^ MMD0 (**Fig. 5h, S10b**), consistent with the suppressed T cell responses observed at 21 dpi. These data associate loss of autophagy gene function with inflammatory and apoptosis pathways in CD11c+ cells, which results in the accumulation of PMN-MDSCs in response to high-dose Mtb infection.

Both activated-PMN-MDSC and PMN-MDSC isolated from tumor-bearing mice robustly suppress T cell responses^15^, suggesting that the PMN-MDSC populations that in the lungs of high-dose Mtb infected *Atg14*^*f/f*^*-LysM-cre* and *Atg14*^*f/f*^*-CD11c-cre* mice at 21 dpi could be responsible for the decreased T cell numbers at this timepoint (**Fig. 3e**). Analysis of the T cell clusters within the scRNA-seq datasets identified 6 sub-clusters, including *Tbx21*+ Th1 cells (T0), *Ccr7*+ naïve T cells (T2, T5), and *Mki67*+ proliferating T cells (T1) (**Fig. 5i**). Upon Mtb infection, T cells skewed towards Th1 associated gene expression, with increased *Tbx21*+ T0 cluster and decreased naïve *Ccr7*+ clusters (**Fig. 5j**). Consistent with the accumulation of PMN-MDSCs, the *Mki67*+ proliferating T cell subcluster (cluster T1, **Fig. 5i**) was reduced in *Atg14*^*f/f*^*-LysM-cre* and *Atg14*^*f/f*^*-CD11c-cre* lungs at 21 dpi (**Fig. 5j**).

Taken together, these results indicate that loss of autophagy in CD11c^+^ cells does not affect early cell intrinsic control of Mtb replication or the initiation of T cell responses during high-dose Mtb infection. Instead, loss of autophagy in CD11c^+^ cells results in elevated accumulation of PMN-MDSCs that harbor high bacterial burden at 21 dpi. The accumulation of PMN-MDSC is associated with decreased T cell proliferation in the lung at 21 dpi, thwarting sustained T cell responses and derailing the control of Mtb infection.

## Acknowledgements

The authors thank Dr. Joel Ernst at UCSF for providing P25/Nur77-GFP/CD45.1 mice, thank Dr. Shumin Tan at Tufts University for sharing pCherry3 plasmid, and Dr. Di Wang at Tsinghua University and Dr. Sairam Andhey at WashU for helpful discussion about scRNA-seq data. The research was supported by NIH grant R01 AI132697, Burroughs Wellcome Fund Investigators in the Pathogenesis of Infectious Disease, and the Philip and Sima Needleman Center for Autophagy Therapeutics and Research (to C.L.S.), NIH grant U19 AI142784 (to C.L.S. and H.W.V.), Tsinghua-Peking Joint Center for Life Sciences (to Y-T.W), and Start-up Foundation of Tsinghua University (to Y-T.W). Authors receive support from Potts Memorial Foundation postdoctoral fellowship (to R.L.K), Stephen I. Morse Fellowship (to S.K.N.), NIH grant T32 AI007172 (to E.M.N.), and NIH grant T32 GM007067 to (S.V.H.). This work was supported, in part, by the Bursky Center for Human Immunology and Immunotherapy Programs at Washington University Immunomonitoring Laboratory. We thank the Core Facility of Center of Biomedical Analysis and Technology Center for Protein Research at Tsinghua University for technical support. We also thank the Genome Technology Access Center at the McDonnell Genome Institute at Washington University School of Medicine. The Center is partially supported by NCI Cancer Center Support Grant #P30 CA91842 to the Siteman Cancer Center and by ICTS/CTSA Grant# UL1TR002345 from the National Center for Research Resources (NCRR), a component of the National Institutes of Health (NIH), and NIH Roadmap for Medical Research. This publication is solely the responsibility of the authors and does not necessarily represent the official view of NCRR or NIH.

## Author contributions

Y-T.W. and C.L.S. designed the project, analyzed the data, and wrote the manuscript. S.F. performed experiments, analyzed the data, analyzed scRNA-seq results, made the final figures, and assisted with manuscript writing. Y-T.W., E.M.N., R.L.K., S.K.N., S.M., N.D., A.S. performed experiments. A.S., S.A., M.N.A. helped design and perform scRNA-seq experiment. S.M.C., X.C., S.V.H., R.W., D.K., and A.S. assisted with experiments and data analysis. H.W.V advised project design. All authors read and edited the manuscript.

## MATERIALS AND METHODS

### Mice

All flox mice (*Atg5*^*f/f* 41^, *Atg7*^*f/f* 42^, *Atg16l1*^*f/f* 43^, *Atg14*^*f/f* 18,44^, *Fip200*^*f/f*^, *Becn1*^*f/f*^) used in this study have been described previously and colonies are maintained in an enhanced barrier facility^5,17^. LysM-cre (Jax #004781), CD11c-cre (Jax #007567), Mrp8-cre (Jax #021614) from the Jackson Laboratory were crossed to specific flox mice. P25/Nur77-GFP/CD45.1 mice^26^ were a gift from Dr. Joel Ernst. Male and female littermates (aged 8–12 weeks) were used and were subject to randomization. Statistical consideration was not used to determine mouse sample sizes. The mice were housed and bred at Washington University in St. Louis and Tsinghua University in specific pathogen-free conditions in accordance with federal and university guidelines, and protocols were approved by the Animal Studies Committee of Washington University and Tsinghua University.

### *M. tuberculosis* infection in mice

*Mycobacterium tuberculosis* Erdman, GFP-Mtb Erdman^45^, and mCherry-expressing Mtb Erdman (mCherry-Mtb) were used in all experiments. mCherry-Mtb was generated by transforming the wild-type strain Erdman with the pCherry3 plasmid^46,47^. Mtb was cultured at 37°C in 7H9 (broth) or 7H11 (agar) (Difco) medium supplemented with 10% oleic acid/albumin/dextrose/catalase (OADC), 0.5% glycerol, and 0.05% Tween 80 (broth). Cultures of GFP-Mtb and mCherry-Mtb were grown in the presence of kanamycin or hygromycin, respectively, to ensure plasmid retention. Mtb cultures in logarithmic growth phase (OD600 = 0.5–0.8) were washed with PBS + 0.05% Tween-80, sonicated to disperse clumps, and diluted in sterile water before delivering 1000 CFUs of aerosolized Mtb per lung using an Inhalation Exposure System (Glas-Col). Within 2 hours of each infection, lungs were harvested from at least two control mice, homogenized, and plated on 7H11 agar to determine the input CFU dose. At each time point after infection, Mtb titers were determined by homogenizing the superior, middle, and inferior lobes of the right lung and plating serial dilutions on 7H11 agar. Colonies were counted after 3 weeks of incubation at 37°C in 5% CO2.

### T cell transfer and neutrophil depletion

For tracing Mtb specific T cells, 4-5 ×10^6^ P25 TCR-Tg CD4^+^ T cells (CD45.1) were labeled with CellTrace (Invitrogen) and adoptively transferred by tail vein injection in 100 µL of sterile PBS 24 hours (h) before mice were infected by Mtb. For neutrophil depletion, mice were intraperitoneally injected every 48 h with 200 µg monoclonal anti-Ly6G antibody (clone 1A8; Leinco) or 200 µg polyclonal rat serum IgG (Sigma-Aldrich) diluted in sterile PBS beginning at 8 dpi and ending at 28 dpi.

### Flow cytometry and cytokine analysis

Mouse lungs (left lobe) were excised after PBS perfusion, placed in DMEM containing 10% FBS, minced finely, and digested at 37°C for an hour with mechanical disruption with a stir bar and enzymatic digestion with Liberase Blendzyme III (Roche), hyaluronidase (Sigma), and DNase I (Sigma). Mediastinal lymph nodes were digested with collagenase B and DNase I (Sigma). Digested tissues were treated with ACK buffer to remove red blood cells and passed through a 70 μm cell strainer to generate single cell suspensions. Cells were stained with Zombie-violet or Zombie-NIR before resuspending in PBS with 2mM EDTA, 3% FBS, and anti-FcγRII/III for blocking. Cells were then labeled with specific antibodies against CD45, Ly6G, CD11c, CD64, MerTK, Ly6C, CD11b, I-A/I-E, siglecF, Gr-1, TCRb, CD4, CD8a, CD62L, CD44 (Biolegend or BD Biosciences). Flow cytometric analysis was performed on an LSRFortessa (BD Biosciences) and Aurora (Cytek), and data was analyzed with FlowJo software (Tree Star). Total cell number was multiplied by the percentage of specific cell type in total single cells, as analyzed by flow cytometry.

For cytokine analysis, lung homogenates were filter sterilized and then analyzed with Mouse Chemokine/Cytokine Panel 1A (ThermoFisher) on a Luminex xMAP FlexMAP3D (Luminex Corp) with Millipore Sigma Belysa software.

### Single-cell RNAseq

scRNA-seq sample preparation was done according to the manufacturer instructions (10x Genomics). Briefly, single cells from naïve or infected lungs were enriched for live cells using dead-cell depletion kit (Miltenyi). Then single cell suspensions were subjected to droplet-based massively parallel scRNA-seq using the Chromium Single Cell 3′ (v3) Reagent Kit in the BSL-3 laboratory as per manufacturer’s instructions (10x Genomics). Cell suspensions were loaded at 1,000 cells/μl to capture 10,000 cells/lane. The 10x Chromium Controller generated GEM droplets, where each cell was labeled with a specific barcode, and each transcript was labeled with a unique molecular identifier (UMI) during reverse transcription. The barcoded cDNA was isolated and removed from the BSL-3 space for library generation. The cDNA underwent 11 cycles of amplification, followed by fragmentation, end repair, A-tailing, adaptor ligation, and sample index PCR as per the manufacturer’s instructions. Libraries were sequenced on a NovaSeq S4 (Illumina), targeting 50,000 read pairs/cell.

The Cell Ranger Single-Cell Software 3.0 available on the 10x Genomics website was used to perform sample demultiplexing. We aligned the resulting fastq files on mouse genome (Ensembl 98) with Cellranger counts. Cellranger counts were processed through the R package Seurat 4^48^. We filtered cells that (1) had more than 5% of mitochondrial gene content, (2) had less than 500 detected genes. Data was log-normalized with a scale factor of 10,000. Dimensionality reduction and clustering were done by detecting the most variable genes using the FindVariableFeatures function. Latent variables (number of UMI’s and mitochondrial content) were regressed out using a negative binomial model (function ScaleData). A UMAP dimensionality reduction was performed on the scaled matrix using the first 30 PCA components to obtain a two-dimensional representation of the cell states. For clustering, we used the functions FindClusters (resolution 0.5). To identify marker genes, we used FindAllMarkers to compare each cluster against all other clusters. For each cluster, only genes that were expressed in more than 25% of cells with at least 0.25-logfold differences were considered. Clusters were annotated using SingleR^49^. Non-immune cells that had low CD45 (Ptprc) expression were excluded, and doublets were removed based on scDblFinder^50^. Designated cell types were sub-clustered using Findclusters (resolution 0.3 for PMN, 0.1 for MMD and T cells), and FindMarkers were used to identify differentially expressed genes (DEgenes). DEgenes of PMN1 or PMN4 were defined by comparing these sub-clusters to all PMNs. Pathway analysis with GSEA on DEgenes of neutrophil subclusters PMN1 and PMN4 was done by using cluster Profiler 4.0^51^. Specifically, (HALLMARK, KEGG and REACTOME), as well as 2 custom defined MDSC gene sets (**Fig. S7**) from published dataset (GSE163834) ^34^. Top DEgenes of PMN1 and PMN4 were interrogated in MDSC clusters of published cancer database^34^ to generate dot plots using ggplot2 in R. These DEgenes were also compared to published marker genes in active TB patients and hits were indicated as arrows on dotplot^16,52^. CD11c+ subclusters were chosen from MMD on resolution 0.3 based on log2FC of CD11c (*Itgax*) (Fig. S9). DEgenes of infected *Atg14*^*f/f*^*-LysM-cre* or *Atg14*^*f/f*^*-CD11c-cre* were defined by comparing to infected *Atg14*^*f/f*^ within CD11c+ MMDs or MMD0. Pathway analysis with GSEA of DEgenes was done by using cluster Profiler 4.0, to compare between infected groups.

### Statistical analysis for biological experiments

All data are from at least two independent experiments. Samples represent biological (not technical) replicates of mice randomly sorted into each experimental group. No blinding was performed during animal experiments. Statistical differences were calculated using Prism (9.0; GraphPad Software) using log-rank Mantel-Cox tests (survival), unpaired two-tailed Student’s t tests, or unpaired two-tailed Mann-Whitney tests. Sample sizes were sufficient to detect differences as small as 10% using the statistical methods described. When used, center values and error bars represent means ± s.e.m. *P* < 0.05 was considered significant. *P* >0.05 was denoted *, ** for *P* < 0.01, *** *P* < 0.001, and **** *P* < 0.0001. Data are presented as mean ± s.e.m. Notable comparisons that were not significantly different are designated as not significant (ns).

## Data availability

scRNA-seq data have been deposited in the NCBI Gene Expression Omnibus (GEO) database are accessible through accession number GSE201410. All other relevant data are available from the corresponding author upon reasonable request.

